## Supplementary Information for "Gelatinase regulates the egress of intracellular replicating populations during *Enterococcus faecalis* infection"

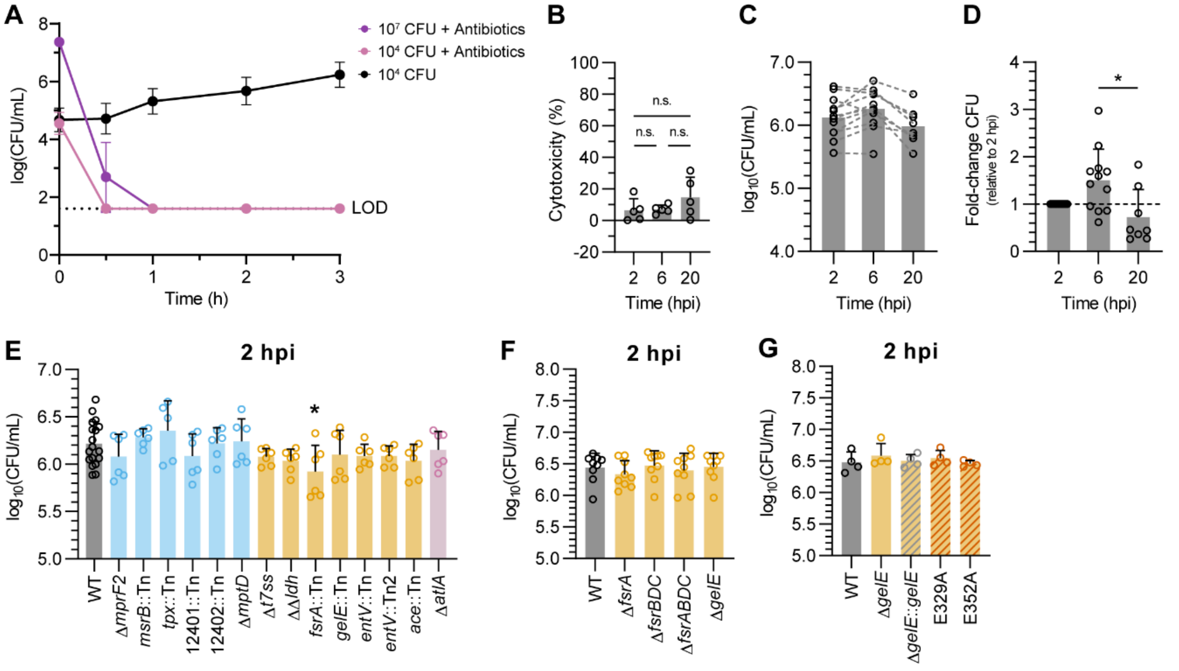


**S1 Fig. Intracellular CFU of *E. faecalis* in RAW264.7 macrophages peaks at 6 hpi for WT OG1RF and is comparable at 2 hpi across OG1RF strains tested. (A)** Antibiotic killing kinetics under conditions used for extracellular killing of *E. faecalis* in the antibiotic protection assay. *E. faecalis* OG1RF log-phase cultures inoculated at $10^7$ (MOI 10 equivalent; purple lines) or $10^4$ CFU (pink lines) are not recovered after ≥ 1 h of vancomycin (10 µg/mL) + gentamicin (150 µg/mL) treatment in DMEM + 10% FBS, in the absence of host cells. Data points are mean ± SD of n = 4. LOD = limit of detection. **(B)** Cytotoxicity measurements of infected macrophages during the antibiotic protection assay. WT OG1RF infection data, shown here for ease of presentation, are also shown in Fig 5A in the experimental context (comparison to mutant strains) they were originally collected from. Statistical significance between timepoints was assessed by one-way ANOVA with Tukey's multiple comparisons test (n = 5). n.s. = not significant. **(C)** Intracellular CFU in RAW264.7 macrophages infected with WT OG1RF at 2, 6, and 20 hpi using the antibiotic protection assay. Counts from the same biological replicate are connected by dotted lines (n = 8-12). **(D)** Fold-change analysis of intracellular CFU quantified in (C), normalised to the intracellular CFU at 2 hpi for each biological replicate. Dotted line indicates baseline CFU at 2 hpi (fold-change CFU = 1.0). (n = 8-12) **(E-G)** Intracellular CFU in RAW264.7 macrophages infected with OG1RF-derived **(E)** mutants of genes implicated in intracellular persistence (blue), virulence (orange), or GelE proteolytic targets (pink), **(F)** genetic deletion mutants of the *fsr* operon and *gelE*, or **(G)** *gelE*-complemented strains at 2 hpi. Statistical significance of each strain against WT was assessed using one-way ANOVA with Dunnett's multiple comparisons test. * = $p < 0.05$.

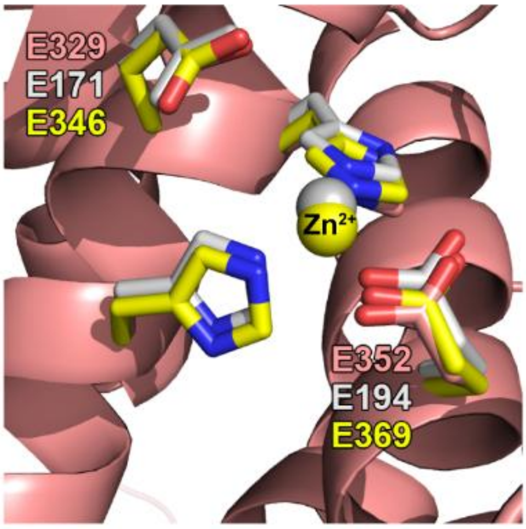


**S2 Fig. Catalytic E329 and zinc-coordinating E352 in the GelE active site are structurally conserved in GelE and other M4-family metalloproteases.** Identification of putative key GelE active site residues E329 and E352 for proteolytic activity, based on structural homology of AlphaFold2-predicted GelE structure (pink) to other M4 family zinc metalloproteases ProA (PDB 6YA1; white) and vibriolysin MCP-02 (PDB 3NQX; yellow). Only mutations in homologous residues E346 and E369 of MCP-02 produced stable, non-proteolytic proteases. Protein structures were aligned in WinCoot v1.1.18 using the Secondary Structure Matching (SSM) Superpose function and visualised in PyMOL v2.5.3.

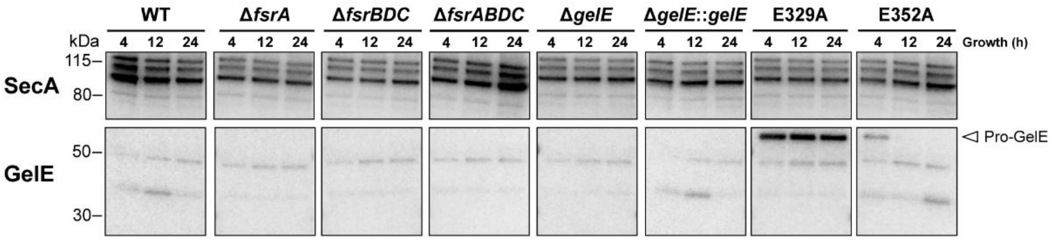


**S3 Fig. Proteolytic mutants E329A and E352A show defects in extracellular secretion and autocatalytic processing of GelE.** Detection of intracellular GelE from *E. faecalis* cell lysates at 4, 12 and 24 h (harvested together with supernatants in **Fig 1G, J**). The membrane protein SecA was included as loading controls. White arrowheads = Pro-GelE (~55 kDa). Images from n = 1 are shown.

**A**

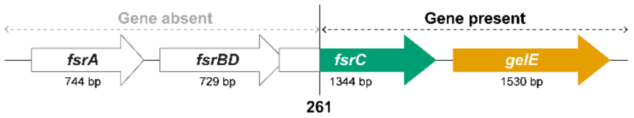


| Strain | ST | *fsrA* | *fsrBDC* | *gelE* | *fsrA* match Start | *fsrA* match End | *fsrBD* match Start | *fsrBD* match End | *fsrC* match Start | *fsrC* match End | *gelE* match Start | *gelE* match End |
|---|---|---|---|---|---|---|---|---|---|---|---|---|
| 01_EF | 179 | – | PARTIAL | + | - | - | - | - | 261 | 1344 | 1 | 1530 |
| 02_EF | 179 | – | PARTIAL | + | - | - | - | - | 261 | 1344 | 1 | 1530 |
| 04_EF | 776 | – | PARTIAL | + | - | - | - | - | 261 | 1344 | 1 | 1530 |
| 06_EF | 179 | – | PARTIAL | + | - | - | - | - | 261 | 1344 | 1 | 1530 |
| 08_EF | 179 | – | PARTIAL | + | - | - | - | - | 261 | 1344 | 1 | 1530 |
| 10_EF | 776* | – | PARTIAL | + | - | - | - | - | 261 | 1344 | 1 | 1530 |
| 12_EF | 776* | – | PARTIAL | + | - | - | - | - | 261 | 1344 | 1 | 1530 |
| 16_EF | 776 | – | PARTIAL | + | - | - | - | - | 261 | 1344 | 1 | 1530 |
| 29_EF | 179 | – | PARTIAL | + | - | - | - | - | 261 | 1344 | 1 | 1530 |
| 31_EF | 179 | – | PARTIAL | + | - | - | - | - | 261 | 1344 | 1 | 1530 |
| 32_EF | 179 | – | PARTIAL | + | - | - | - | - | 261 | 1344 | 1 | 1530 |
| 37_EF | 179 | – | PARTIAL | + | - | - | - | - | 261 | 1344 | 1 | 1530 |
| 41_EF | 64* | – | PARTIAL | + | - | - | - | - | 261 | 1344 | 1 | 1530 |
| EF_1008 | 776 | – | PARTIAL | + | - | - | - | - | 261 | 1344 | 1 | 1530 |
| EFS_M02 | 179 | – | PARTIAL | + | - | - | - | - | 261 | 1344 | 1 | 1530 |
| EFS_M03 | 179 | – | PARTIAL | + | - | - | - | - | 261 | 1344 | 1 | 1530 |
| EFS_M04 | 179 | – | PARTIAL | + | - | - | - | - | 261 | 1344 | 1 | 1530 |

**B**

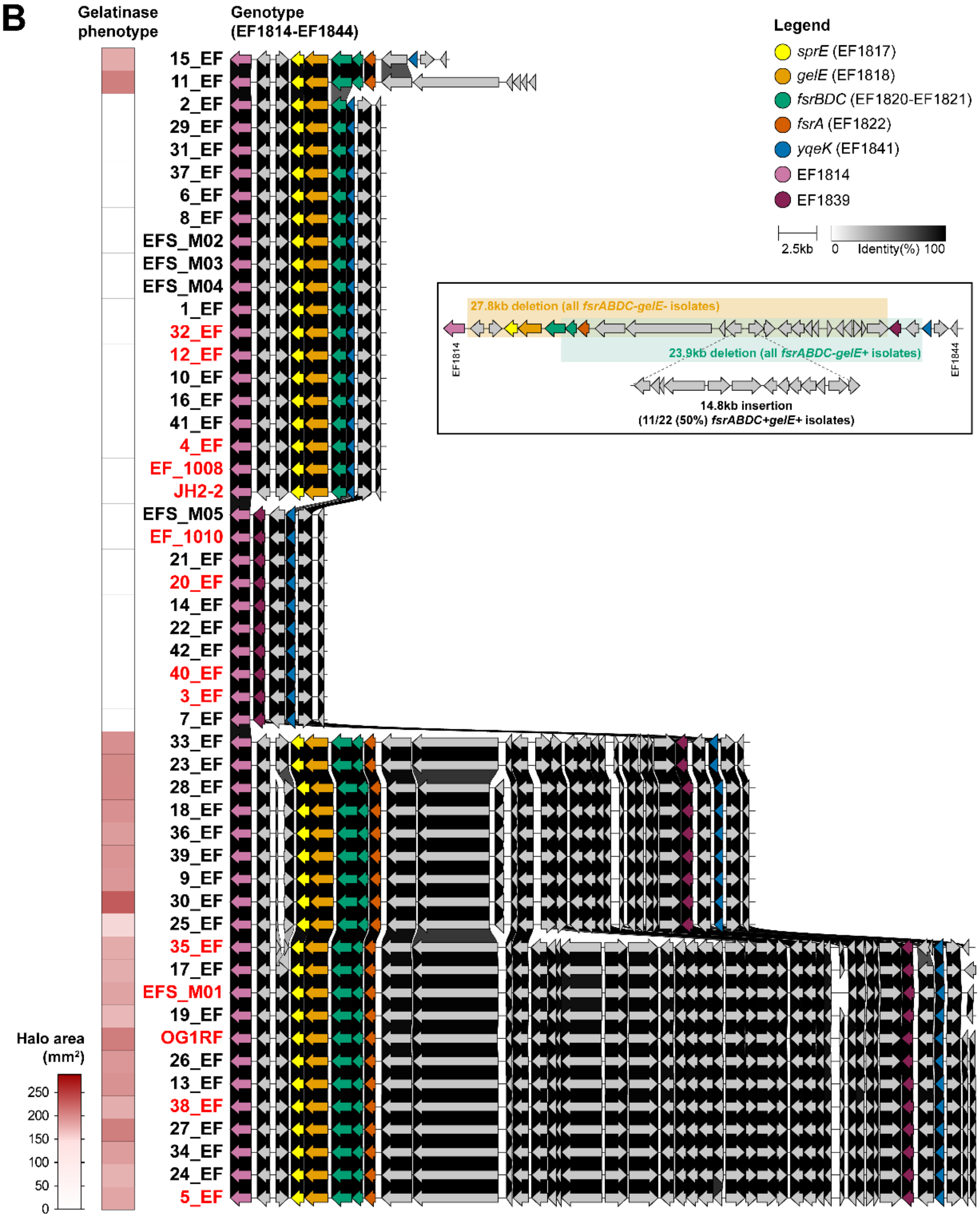

**S4 Fig. *fsrABDC-gelE+* and *fsrABDC-gelE-* wound isolates of *E. faecalis* show distinct and conserved genomic deletions. (A)** Matching blastn query matches of contigs from all *fsrABDC-gelE+* strains to *fsrA, fsrBDC, fsrC or gelE.* The “Start” and “End” columns indicate nucleotide positions of matches in the respective genes, while dashes indicate no matches, as visually represented by the schematic diagram. Sequence type (ST) of each strain is shown, with asterisks (*) indicating uncertainties in SRST2 sequence type calling. **(B)** Alignment of the genomic region corresponding to EF1814-EF1844 of the V583 reference genome for *E. faecalis* wound isolates. OG1RF and JH2-2 were included as reference strains. Genes with sequence identity > 30% are connected by shaded lines, colored by identity (%) as indicated in the legend. Colored arrows indicate *fsrA, fsrBDC, gelE* and *sprE* operons, as well as other genes flanking genomic deletion regions. Inset shows a diagrammatic summary of the conserved genomic deletions or insertions observed. The corresponding GelE producing phenotype for each strain (as reported in **Fig 3A**) are shown on the left. Strains highlighted in red are further tested by *in vitro* antibiotic protection assay (**Fig 3B-D and S5**).

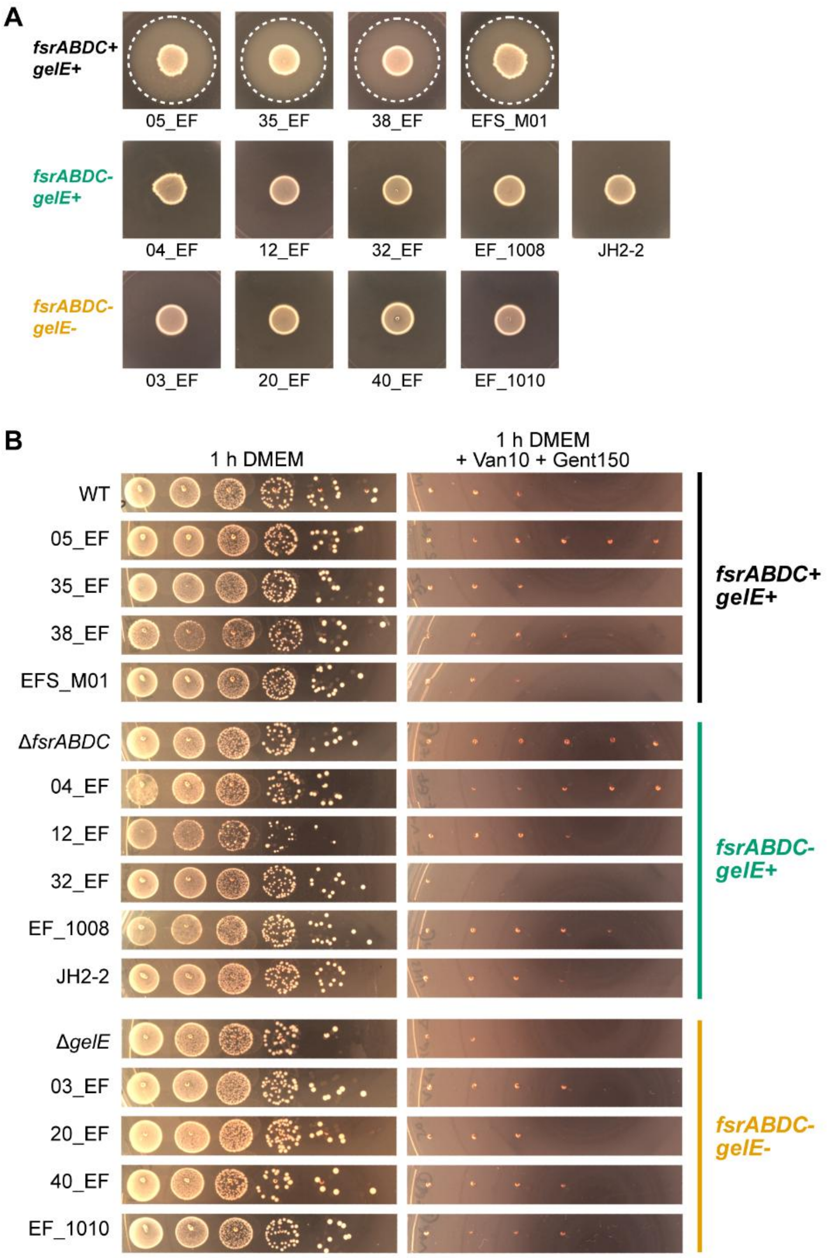


**S5 Fig. Selected wound isolates of *E. faecalis* exhibit gelatinase activity expected from their genotype and are phenotypically gentamicin-susceptible for the antibiotic protection assay. (A)** Gelatinase activity of *E. faecalis* wound isolates and JH2-2 on Todd-Hewitt agar + 3% gelatin at 24 h. Halo formation (white dashed line) indicates gelatinase activity. Genotypes of wound isolates are indicated on the left. Representative images of n = 3 are shown. (B) Validation of bactericidal activity of vancomycin (10 µg/mL) + gentamicin (150 µg/mL) on log-phase cultures of *E. faecalis* wound isolates and laboratory strains (JH2-2; OG1RF WT, Δ*fsrABDC*, Δ*gelE*). Representative images from post-treatment serial dilutions are shown (n = 3). Left column = untreated controls.

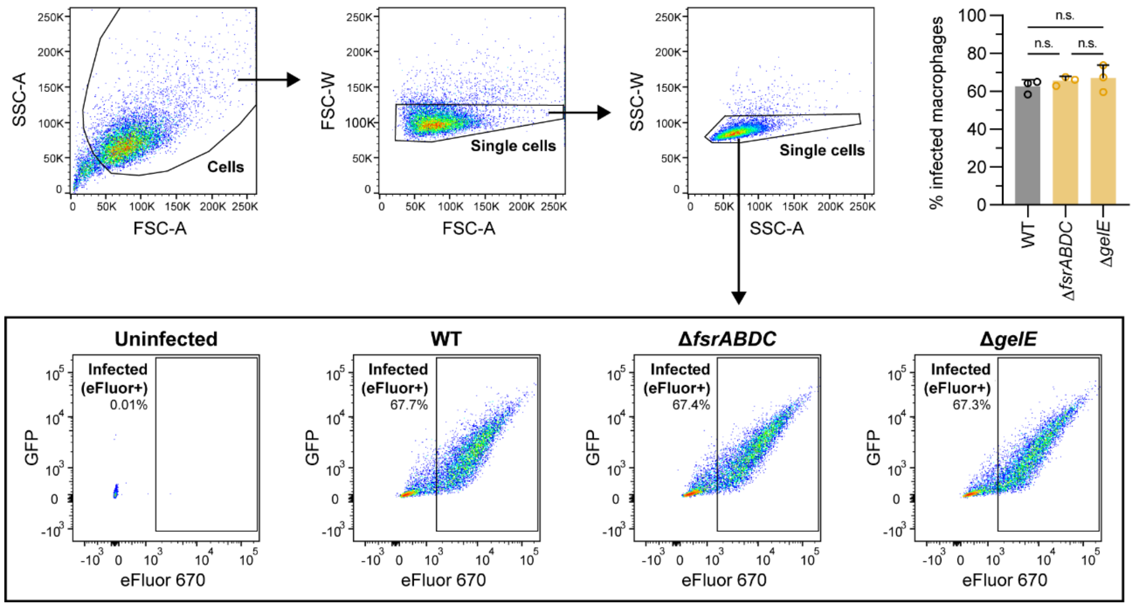


**S6 Fig. Infection efficiency at 2 hpi is comparable between GelE+ WT and GelE- deletion strains.** Flow cytometry analysis of RAW264.7 macrophages at 2 hpi infected with GFP-expressing WT (GelE+, gray bar), Δ*fsrABDC* (GelE-, orange bar) or Δ*gelE* (GelE-, orange bar) pre-stained with eFluor 670, showing the gating strategy and proportion of eFluor 670+ macrophages (infected macrophages). Uninfected macrophages are used as gating controls. Representative dotplots from n = 3 are shown. Statistical significance was assessed by one-way ANOVA with Tukey's multiple comparisons test. n.s. = not significant.

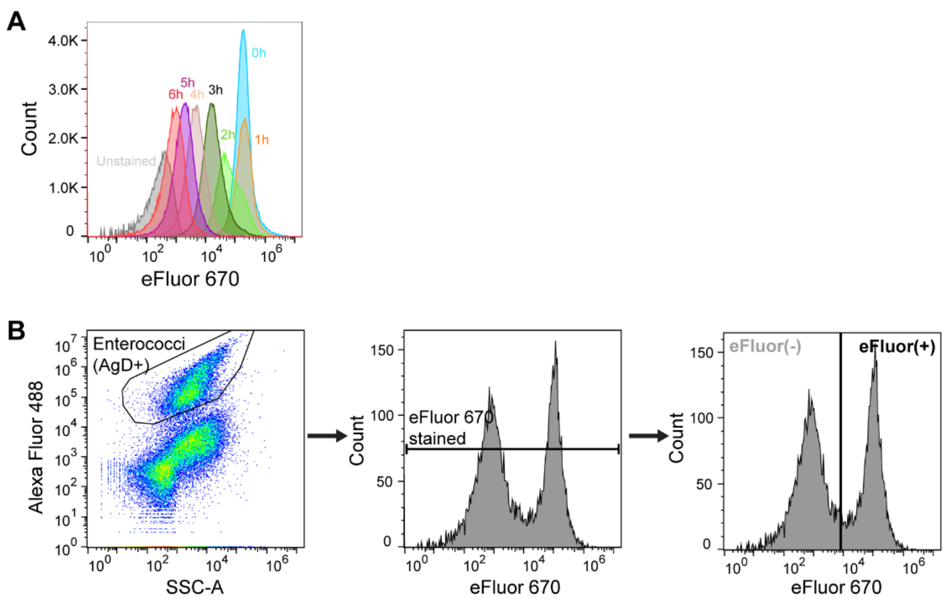


**S7 Fig. Detection of replicating intracellular *E. faecalis* from macrophage lysate by flow cytometry analysis of eFluor 670 proliferation dye.** (A) Serial dilution of eFluor 670 proliferation dye in actively replicating *E. faecalis* pDasherGFP OG1RF cultures in colorless DMEM + 10% FBS over 6 h. Bacterial cultures were aliquoted hourly for flow cytometry analysis. Representative histograms of n = 3 are shown. (B) Gating strategy for eFluor 670 analysis of intracellular *E. faecalis* released from infected RAW264.7 lysates. *E. faecalis* stained for Group D antigen (AgD) was gated as Alexa Fluor 488+ events. Debris at the histogram edges (<$10^0$ and >$10^7$ fluorescence intensity) were gated out prior to histogram bisection for accurate quantification of eFluor$^+$ and eFluor$^-$ events.

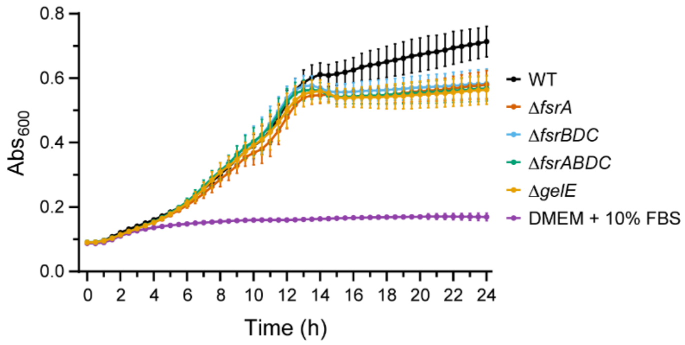


**S8 Fig. *fsr/gelE* deletion mutants of *E. faecalis* OG1RF do not exhibit growth rate differences in macrophage cell culture media.** *fsr/gelE* deletion mutants were grown in colorless DMEM + 10% FBS for 24 h and the absorbance at 600nm (Abs600) was measured at 30 min intervals. Colorless DMEM + 10% FBS is used as a negative control. Growth curve is plotted as a mean ± SD of n = 2 with 10 technical replicates each.

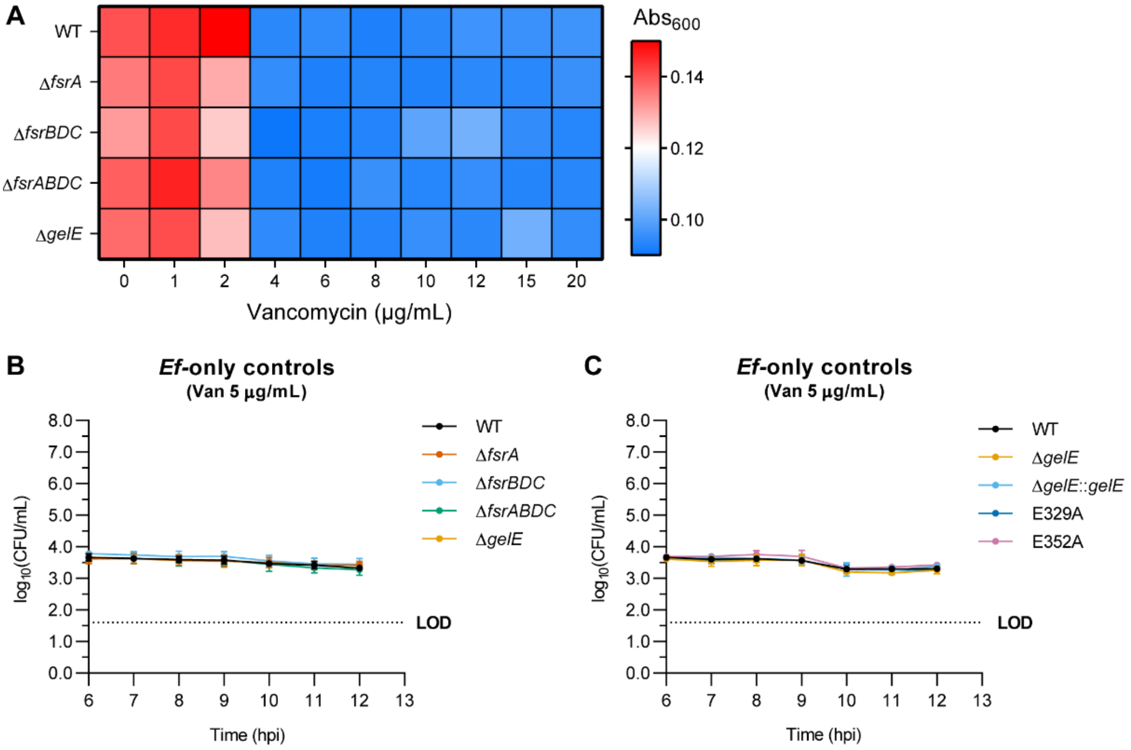


**S9 Fig. 5 µg/mL vancomycin is sufficient for a bacteriostatic effect on extracellular *E. faecalis* with minimal bactericidal effect. (A)** Determination of minimum inhibitory concentration of vancomycin by broth microdilution in colorless DMEM + 10% FBS for OG1RF WT and *fsr/gelE* deletion mutants. Bacterial growth was measured as absorbance at 600 nm ($Abs_{600}$), represented in the heatmap as mean of n = 2. **(B-C)** Validation of bacteriostatic effect of 5 µg/mL vancomycin for **(B)** *fsr/gelE* deletion mutants (n = 4) or **(C)** *gelE*-complemented strains (n = 3). Recovered bacteria CFU is expressed in $log_{10}$(CFU/mL) as mean ± SD of n = 3-4.

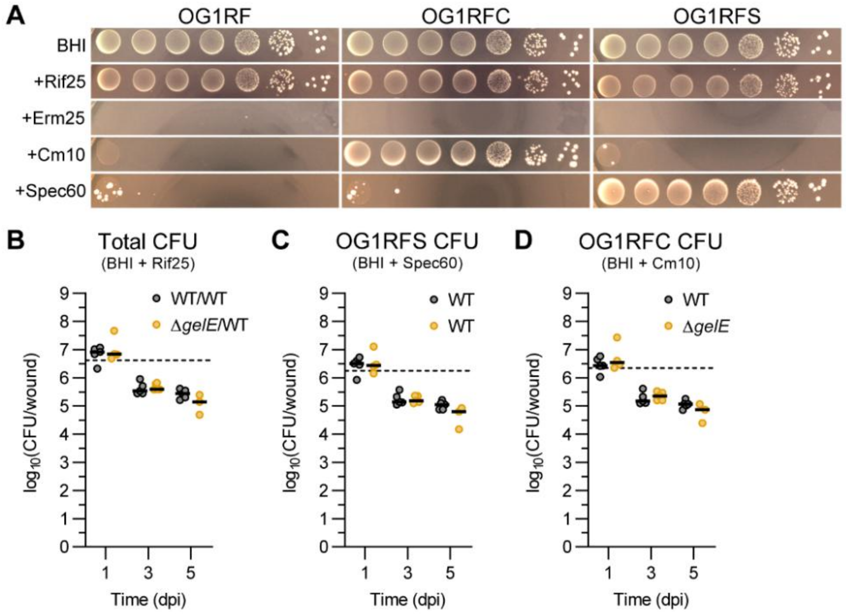


**S10 Fig. Differential CFU quantification of OG1RFC and OG1RFS strains during a competitive model of wound infection.** **(A)** Validation of growth selection of isogenic strains OG1RFC and OG1RFS on chloramphenicol (Cm, 10 µg/mL) and spectinomycin (Spec, 60 µg/mL) BHI plates respectively, compared to parental strain OG1RF. Rifampicin (Rif, 25 µg/mL) and erythromycin (Erm, 25 µg/mL) are used as positive and negative growth controls of antibiotic selection respectively. **(B-D)** CFU quantification from wound homogenates on antibiotic agars, selecting for **(B)** total OG1RFC + OG1RFS, **(C)** OG1RFC, and **(D)** OG1RFS. Black bars represent median of 3-5 animals per infection group from one independent experiment. Statistical significance between infection groups of the same timepoint was assessed using Mann-Whitney test. Only comparisons with $p < 0.05$ are annotated. Dotted lines show bacteria inoculum CFU.

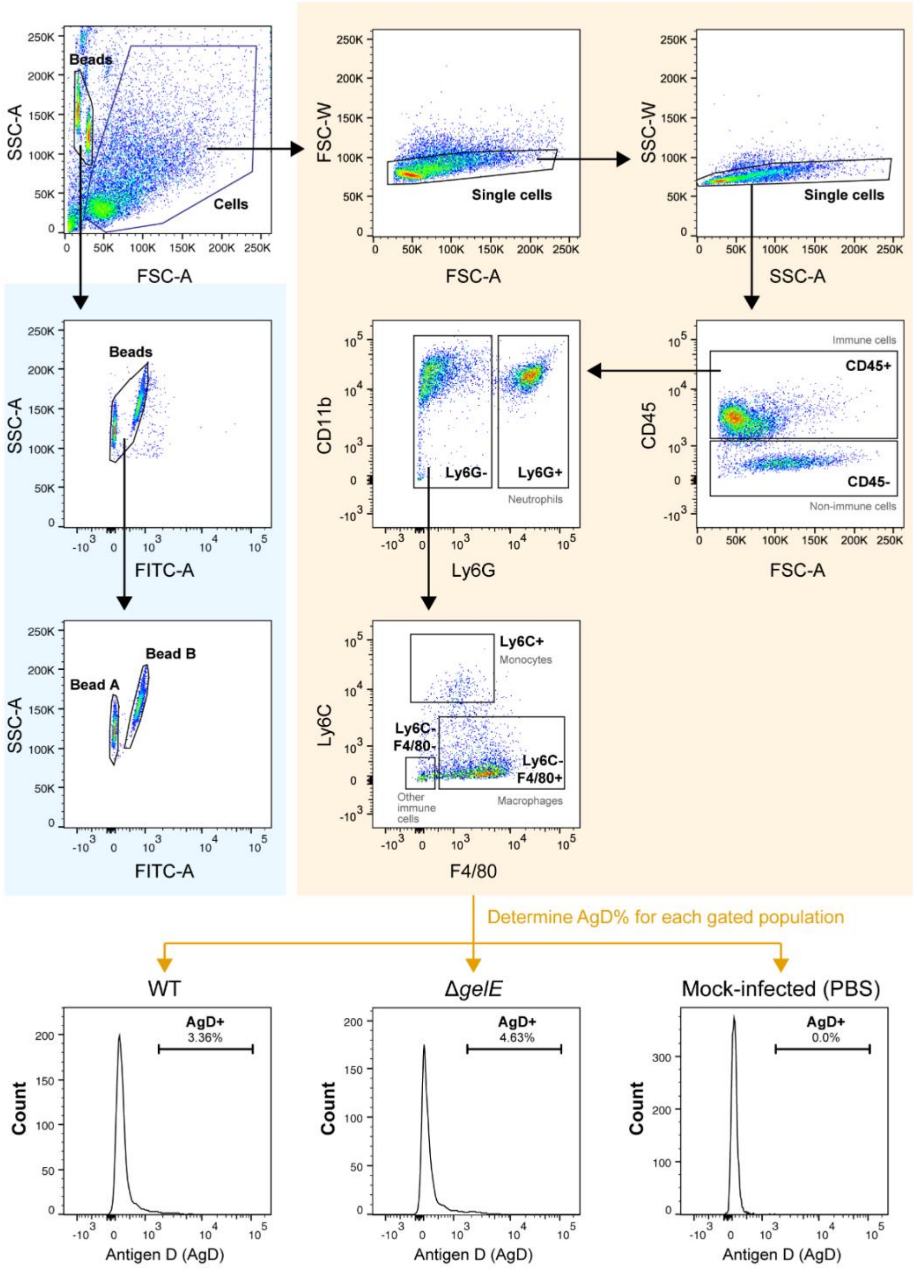


**S11 Fig. Flow cytometry gating and quantification of dissociated cells from 5 dpi WT, Δ*gelE* or mock-infected murine wounds.** Wound cells (yellow panel) as well as AccuCheck Counting Beads A and Beads B (blue panel) were gated from forward-scatter/side-scatter (FSC/SSC) dotplots. Single cells from the wound were then separately gated for CD45+ (immune cells) or CD45- (non-immune cells), and CD45+ cells were further separated into Ly6G+ (neutrophils) and Ly6G- populations. CD45+ Ly6G- populations were subsequently gated into Ly6C+ (monocytes), Ly6C- F4/80+ (macrophages), and Ly6C- F4/80- (other immune cells) populations. Cell counts from each gated population were normalized to the total Bead A + Bead B count from each sample. Each immune cell subpopulation was further analysed into AgD+ (infected) and AgD- (uninfected) cells, with AgD+ threshold determined in histograms based on AgD- mock-infected samples.

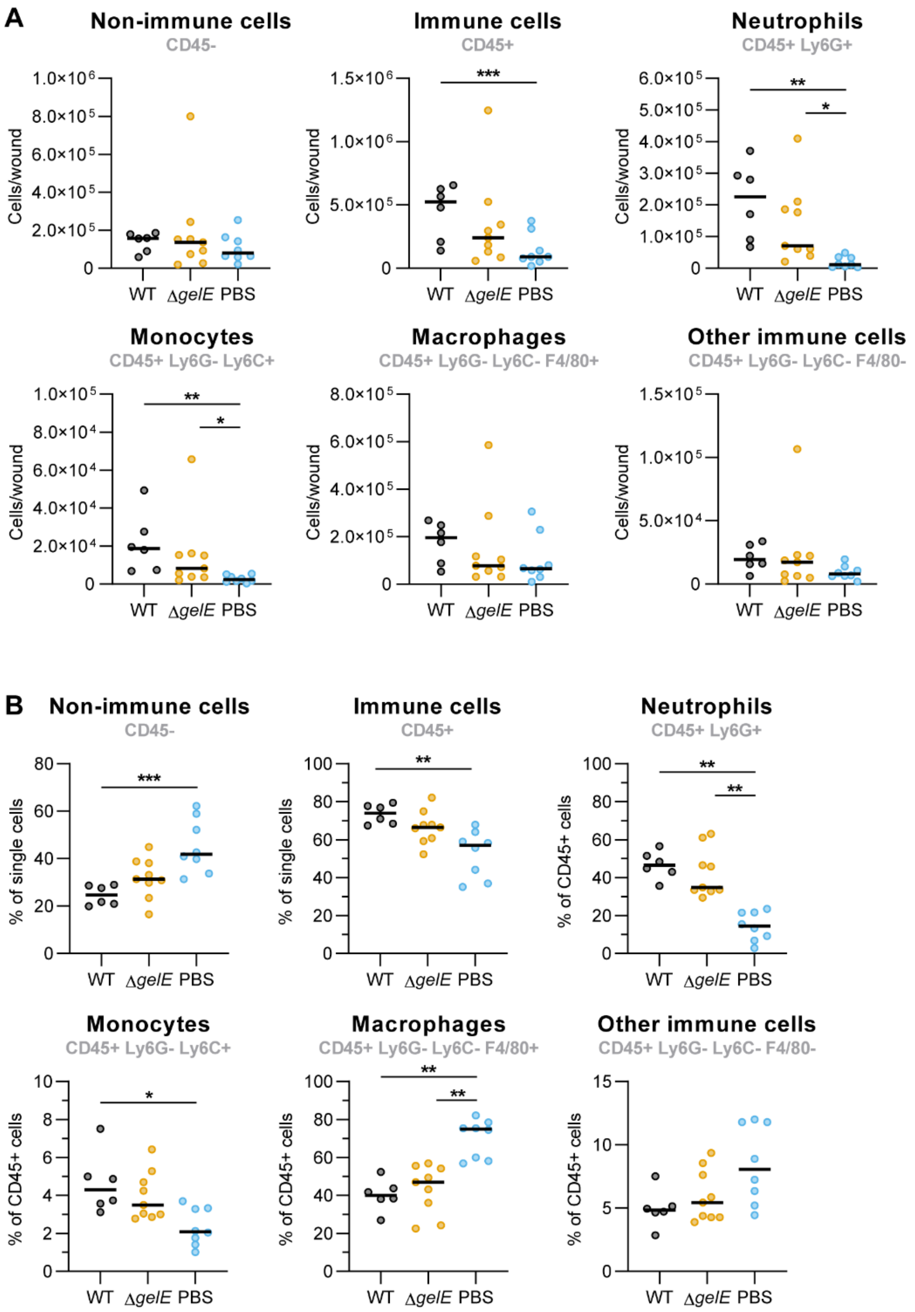


**S12 Fig. Innate immune cell profile of 5 dpi WT, Δ*gelE* or mock-infected murine wounds. (A)** Cell counts from gated populations described in **S11 Fig**, normalized to AccuCheck Counting Bead counts. **(B)** Percentage of gated populations relative to all single cells (CD45+/CD45- population) or all CD45+ cells (immune cell populations). Bars represent median from 6-9 mice from two independent experiments. For each cell population, statistical significance was assessed using Kruskal-Wallis test with Dunn's multiple comparison test. * = $p < 0.05$, ** = $p < 0.01$, *** = $p < 0.001$.

**S1 Table. Strains used in this study.** Numbers after the gene name indicate position (in bp) within the protein coding sequence where the transposon is inserted for gene disruption.

| **Strain name** | **Description** | **Antibiotic resistance*** | **Source** |
|---|---|---|---|
| OG1RF; Wildtype (WT) | Plasmid-free, gelatinase-positive strain with spontaneous Rif and Fus mutations | Rif, Fus | (1) |
| OG1RFC | OG1RF WT with a chromosomal insertion of *cat* (chloramphenicol resistance cassette) at the genomic insertion site for expression (GISE) | Rif, Fus, Cm | This study |
| OG1RFS | OG1RF WT with a chromosomal insertion of *spc* (spectinomycin resistance cassette) at the genomic insertion site for expression (GISE) | Rif, Fus, Spec | This study |
| JH2-2 | Gelatinase-negative strain with spontaneous Rif and Fus mutations | Rif, Fus | (2) |
| Δ*fsrA* | ΔOG1RF_11526 | Rif, Fus | This study |
| Δ*fsrBDC* | ΔOG1RF_11527-11528 | Rif, Fus | This study |
| Δ*fsrABDC* | ΔOG1RF_11526-11528 | Rif, Fus | (3) |
| Δ*gelE* | ΔOG1RF_11529 | Rif, Fus | (4) |
| Δ*gelE*::*gelE* | Δ*gelE* with chromosomal complementation of WT *gelE* sequence with a silent A29A mutation | Rif, Fus | This study |
| E352A | Δ*gelE*::*gelE*$^{E352A}$; Δ*gelE* with chromosomal complementation of *gelE* containing a silent A29A mutation and a zinc-coordinating E352A active site mutation | Rif, Fus | (3) |
| E329A | Δ*gelE*::*gelE*$^{E329A}$; Δ*gelE* with chromosomal complementation of *gelE* containing a silent A29A mutation and a catalytic E329A active site mutation | Rif, Fus | This study |
| Δ*mprF2* | ΔOG1RF_10760 | Rif, Fus | (5) |
| *msrB*::Tn | *msrB*112::EfaMarTn | Rif, Fus, Cm | (6) |
| 12401::Tn | OG1RF_12401(214)::EfaMarTn; part of phosphotransferase system PTS8 | Rif, Fus, Cm | (6) |
| 12402::Tn | OG1RF_12402(190)::EfaMarTn; part of phosphotransferase system PTS8 | Rif, Fus, Cm | (6) |
| *tpx*::Tn | *tpx*349::EfaMarTn | Rif, Fus, Cm | (6) |
| Δ*mptD* | ΔOG1RF_10021; part of phosphotransferase system PTS1 | Rif, Fus | (7) |

| | | | |
|---|---|---|---|
| Δ*t7ss* | ΔOG1RF_11100-OG1RF_11127 | Rif, Fus | (8) |
| ΔΔ*ldh* | Δ*ldh1*Δ*ldh2*; ΔOG1RF_10199ΔOG1RF_10373 | Rif, Fus | (9) |
| *fsrA::*Tn | *fsrA*10*::*EfaMarTn | Rif, Fus, Cm | (6) |
| *gelE::*Tn | *gelE*968*::*EfaMarTn | Rif, Fus, Cm | (6) |
| *sprE*::Tn | *sprE*674::EfaMarTn | Rif, Fus, Cm | (6) |
| *entV*::Tn | *entV*(-5)::EfaMarTn | Rif, Fus, Cm | (6) |
| *entV*::Tn2 | *entV*182::EfaMarTn | Rif, Fus, Cm | (6) |
| *ace*::Tn | *ace*665::EfaMarTn | Rif, Fus, Cm | (6) |
| Δ*atlA* | ΔOG1RF_10533 (EF0799) | Rif, Fus | (10) |
| OG1RFC Δ*gelE* | OG1RFC ΔOG1RF_11529 | Rif, Fus | This study |

*Rif, rifampicin; Fus, fusidic acid; Cm, chloramphenicol.

**S2 Table. Details of the wound clinical isolates tested in this study.** Asterisks (*) indicating uncertainties in sequence type (ST) calling by SRST2. + and – indicates genotype/phenotype presence and absence respectively.

| Strain name | Collection site | ST | *fsrABDC* genotype | *gelE* genotype | GelE activity phenotype |
|---|---|---|---|---|---|
| 1_EF | Left foot | 179 | - | + | - |
| 2_EF | Perianal abscess | 179 | - | + | - |
| 3_EF | Ear cyst wound swab | 16 | - | - | - |
| 4_EF | Scrotal abscess swab | 776 | - | + | - |
| 5_EF | Right foot wound swab | 81 | + | + | + |
| 6_EF | Peripancreatic fluid | 179 | - | + | - |
| 7_EF | Wound swab | 16 | - | - | - |
| 8_EF | Left foot tissue | 179 | - | + | - |
| 9_EF | Suprapubic wound | 6 | + | + | + |
| 10_EF | Nasopharyngeal wound | 776* | - | + | - |
| 11_EF | Right lateral foot | 21 | + | + | + |
| 12_EF | Supraorbital wound | 776* | - | + | - |
| 13_EF | Buttock wound | 116 | + | + | + |
| 14_EF | Forefoot tissue | 16 | - | - | - |
| 15_EF | Left periauricular abscess | 856* | + | + | + |
| 16_EF | Wound swab | 776 | - | + | - |
| 17_EF | Sacral sore | 116 | + | + | + |
| 18_EF | Sacral swab | 6 | + | + | + |
| 19_EF | Index finger | 862 | + | + | + |
| 20_EF | Toe | 16 | - | - | - |
| 21_EF | Bile wound swab | 16 | - | - | - |
| 22_EF | Wound swab | 16 | - | - | - |
| 23_EF | Wound swab | 6 | + | + | + |
| 24_EF | Perianal abscess | 81 | + | + | + |
| 25_EF | Wound swab | 6 | + | + | + |
| 26_EF | Toe wound | 202 | + | + | + |
| 27_EF | Bile | 81 | + | + | + |
| 28_EF | Leg wound | 6 | + | + | + |
| 29_EF | Pancreatic abscess | 179 | - | + | - |
| 30_EF | Sacral swab | 6 | + | + | + |

| | | | | | |
|---|---|---|---|---|---|
| 31_EF | Foot swab | 179 | - | + | - |
| 32_EF | Stump | 179 | - | + | - |
| 33_EF | Wound swab | 6 | + | + | + |
| 34_EF | Heel wound swab | 81 | + | + | + |
| 35_EF | Leg wound | 314 | + | + | + |
| 36_EF | Gluteal pus | 6 | + | + | + |
| 37_EF | Peripancreatic fluid | 179 | - | + | - |
| 38_EF | Inguinal abscess | 81* | + | + | + |
| 39_EF | Tissue right forefoot | 6 | + | + | + |
| 40_EF | Wound swab | 16 | - | - | - |
| 41_EF | Tissue left lateral leg | 64* | - | + | - |
| 42_EF | Wound swab bedsore | 16 | - | - | - |
| EF_1008 | Wound swab | 776 | - | + | - |
| EF_1010 | Wound swab | 16 | - | - | - |
| EFS_M01 | Wound swab | 482 | + | + | + |
| EFS_M02 | Wound swab | 179 | - | + | - |
| EFS_M03 | Wound swab | 179 | - | + | - |
| EFS_M04 | Wound swab | 179 | - | + | - |
| EFS_M05 | Wound swab | 16 | - | - | - |

**S3 Table. Plasmids used in this study.**

| Plasmid name | Description | Antibiotic resistance* | Source |
|---|---|---|---|
| pGCP213 | Temperature-sensitive shuttle plasmid for allelic exchange | Erm | (11) |
| pGCP213::*fsrA_del* | pGCP213::US$_{fsrA}$-DS$_{fsrA}$ | Erm | This study |
| pGCP213::*fsrBDC_del* | pGCP213::US$_{fsrBDC}$-DS$_{fsrBDC}$ | Erm | This study |
| pGCP213::*gelE* | pGCP213::US$_{gelE}$-*gelE*(A29A)-DS$_{gelE}$ | Erm | This study |
| pGCP213::*gelE*$^{E329A}$ | pGCP213::US$_{gelE}$-*gelE*(A29A, E329A)-DS$_{gelE}$ | Erm | This study |
| pGCP213::*gelE*$^{E352A}$ | pGCP213::US$_{gelE}$-*gelE*(A29A, E352A)-DS$_{gelE}$ | Erm | (3) |
| pSD15 | pCR8/GW/TOPO with OG1RF_11779-MCS-OG1RF_11778 (11779-MCS-11778) for GISE constructs | Spec | (12) |
| pBSU101::DasherGFP | pBSU101 with constitutively expressed Dasher GFP under a CFB promoter (P$_{CFB}$) | Spec | (13) |
| pSD15::DasherGFP | pSD15::11779-P$_{CFB}$-DasherGFP-11778 | Spec | This study |
| pSD15::P$_{gelE}$-DasherGFP | pSD15::11779-P$_{gelE}$-DasherGFP-11778 | Spec | This study |
| pRV1 | Replicon-free shuttle plasmid containing P-*pheS** negative selection cassette | Erm | (12) |
| pFR212 | pGCP213::11779-P$_{gelE}$-DasherGFP-11778 | Erm | This study |
| pFR213 | pGCP213-P-*pheS**::11779-P$_{gelE}$-DasherGFP-11778 | Erm | This study |
| pFR212::*cat* | pGCP213::11779-*cat*-11778 | Erm, Cm | This study |
| pFR212::*spc* | pGCP213::11779-*spc*-11778 | Erm, Spec | This study |
| pFR213::*gelE*_del | pFR213::US$_{gelE}$-DS$_{gelE}$ | Erm | This study |

*Erm, erythromycin; Spec, spectinomycin; Cm, chloramphenicol.

**S4 Table. Primers used in this study.**

| Purpose | Primer name | Sequence | Description |
|---|---|---|---|
| Creation of Δ*fsrA* | oFR1 | GATGCATGCTCGAGCCATCAATGATCAAACGATTTCGTA | Primer 1, for amplifying 500 bp upstream sequence of *fsrA* from WT OG1RF. 15 bp overlap for InFusion cloning is underlined. |
| | oFR2 | CTTAATTAGTCATTATCCCTCCCTAAGTAGCCATTTGTTCACTCATCC | Primer 2, for amplifying 500 bp upstream sequence of *fsrA* from WT OG1RF and ligation to downstream sequence. Complementary to oFR3. |
| | oFR3 | GGATGAGTGAACAAATGGCTACTTAGGGAGGGATAATGACTAATTAAG | Primer 3, for amplifying 500 bp downstream sequence of *fsrA* from WT OG1RF and ligation to upstream sequence. Complementary to oFR2. |
| | oFR4 | TACCGAGCTCGGATCCGTTTAAATCAGATGGCTGAACAG | Primer 4, for amplifying 500 bp downstream sequence of *fsrA* from WT OG1RF. 15 bp overlap for InFusion cloning is underlined. |
| | oFR7 | GCTCGAGCATGCATCTAGAGG | Primer F, for inverse PCR (linearisation) of pGCP213 to insert *fsrA* deletion cassette. 15 bp overlap for InFusion cloning is underlined. |
| | oFR8 | GATCCGAGCTCGGTACCAAG | Primer R, for inverse PCR (linearisation) of pGCP213 to insert *fsrA* deletion cassette. 15 bp overlap for InFusion cloning is underlined. |
| | M13F_ pUC (-40) | GTTTTCCCAGTCACGAC | Universal sequencing primer F for pFR213 inserts. |
| | M13R_ pUC (-26) | GTCATAGCTGTTTCCTG | Universal sequencing primer R for pFR213 inserts. |
| | oFR5 | TGACAAGAACAGTTTGGCGG | Sequencing primer F for *fsrA* deletion |
| | oFR6 | AAGGTTTCGCTTAACGTCCC | Sequencing primer R for *fsrA* deletion |
| Creation of Δ*fsrBDC* | oFR15 | GATGCATGCTCGAGCCAATTTATTCAAAAGAACGTGGATTTTC | Primer 1, for amplifying 502 bp upstream sequence of *fsrBDC* from WT OG1RF. 15 bp overlap for InFusion cloning is underlined. |
| | oFR16 | CCATAGCAAAAAAGTTGTTAACAAATTCATTCCATATCGCCCTCCTCTTCAAG | Primer 2, for amplifying 502 bp upstream sequence of *fsrBDC* from WT OG1RF and ligation to downstream sequence. Complementary to oFR3. |
| | oFR17 | CTTGAAGAGGAGGGCGATATGGAATGAATTTGTTAACAACTTTTTTGCTATGG | Primer 3, for amplifying 507 bp downstream sequence of *fsrBDC* from WT OG1RF and ligation to upstream sequence. Complementary to oFR2. |
| | oFR18 | TACCGAGCTCGGATCCACATTATCTGAATCAACAGTAACGC | Primer 4, for amplifying 507 bp downstream sequence of *fsrBDC* from WT OG1RF. 15 bp overlap for InFusion cloning is underlined. |
| | oFR7 | GCTCGAGCATGCATCTAGAGG | Primer F, for inverse PCR (linearisation) of pGCP213 to insert *fsrBDC* deletion cassette. |

| | | | |
|---|---|---|---|
| | | | 15 bp overlap for InFusion cloning is underlined. |
| | oFR8 | GATCCGAGCTCGGTACCAAG | Primer R, for inverse PCR (linearisation) of pGCP213 to insert *fsrBDC* deletion cassette. 15 bp overlap for InFusion cloning is underlined. |
| | M13F_ pUC (-40) | GTTTTCCCAGTCACGAC | Universal sequencing primer F for pFR213 inserts. |
| | M13R_ pUC (-26) | GTCATAGCTGTTTCCTG | Universal sequencing primer R for pFR213 inserts. |
| | oFR19 | ACGGAACGGAAGTACCAGTC | Sequencing primer F for *fsrBDC* deletion |
| | oFR20 | AGCTGCCTCAGAAATTGCCT | Sequencing primer R for *fsrBDC* deletion |
| Creation of *gelE*-complementation strains | oFR94 | GATGCATGCTCGAGCGAATTGAAAATGTTCGCTATCTC | Primer 1, for amplifying *gelE* from WT OG1RF (including 519 bp upstream sequence). 15 bp overlap for InFusion cloning is underlined. |
| | oFR96 | GAATAAACTTGTTCTTC**CGC**GGC | Primer 2, mutagenic primer for A29A silent mutation. Overlaps with oFR99 (primer 3). The GCA > GCG silent mutation is in bold. |
| | oFR99 | GTAGCC**GCG**GAAGAACAAG | Primer 3, mutagenic primer for A29A silent mutation. Overlaps with oFR96 (primer 2). The GCA > GCG silent mutation is in bold. |
| | oFR98 | TACCGAGCTCGGATCGACGATCGTTTTGTTTGC | Primer 4, for amplifying *gelE* from WT OG1RF (including 526 bp downstream sequence). 15 bp overlap for InFusion cloning is underlined. |
| | oFR80 | GTTGGTCAT**GCA**ATGACACATGGTG | Mutagenic primer 1 for E329A site-directed mutagenesis of *gelE*. Overlaps with oFR81. The GAA > GCA (E > A) mutation is in bold. |
| | oFR81 | CACCATGTGTCAT**TGC**ATGACCAAC | Mutagenic primer 2 for E329A site-directed mutagenesis of *gelE*. Overlaps with oFR80. The GAA > GCA (E > A) mutation is in bold. |
| | oFR82 | CAGGTGCCTTGAAT**GCA**TCTTATTCTG | Mutagenic primer 1 for E352A site-directed mutagenesis of *gelE*. Overlaps with oFR83. The GAA > GCA (E > A) mutation is in bold. |
| | oFR83 | CAGAATAAGA**TGC**ATTCAAGGCACC | Mutagenic primer 2 for E352A site-directed mutagenesis of *gelE*. Overlaps with oFR82. The GAA > GCA (E > A) mutation is in bold. |
| | M13F_ pUC (-40) | GTTTTCCCAGTCACGAC | Universal sequencing primer F for pGCP213 inserts. |
| | M13R_ pUC (-26) | GTCATAGCTGTTTCCTG | Universal sequencing primer R for pGCP213 inserts. |
| | oFR109 | GCAACAAATATTTACGCAGGGAAAG | Sequencing primer F for colony PCR of correct *gelE*-complemented clones. |

| | | | |
|---|---|---|---|
| | oFR110 | GACAGGCTAAAACCGGCTATC | Sequencing primer R for colony PCR of correct *gelE*-complemented clones. |
| Construction of pFR212/ pFR213 | oFR43 | GGCGGCCGCGAGCTCAATTCATCTAAAATAGTACGCTTCT | Primer F, for amplifying $P_{CFB}$-DasherGFP insert from pBSU101::DasherGFP. SacI restriction site is underlined. |
| | oFR44 | GGGTACCACGCATGCTTACTGATACGTGTCCAGATC | Primer R, for amplifying $P_{CFB}$-DasherGFP insert from pBSU101::DasherGFP. SphI restriction site is underlined. |
| | oFR102 | CTATTTTAGATGAATTGAGCTCGCGG | Primer F, for inverse PCR (linearisation) of pSD15::DasherGFP for $P_{gelE}$ replacement. 15 bp overlap for InFusion cloning is underlined. |
| | oFR103 | ATGACGGCATTGACGGAAG | Primer F, for inverse PCR (linearisation) of pSD15::DasherGFP for $P_{gelE}$ replacement. 15 bp overlap for InFusion cloning is underlined. |
| | oFR104 | ATTCATCTAAAATAGGAGTTATGAGGGGCAATACAG | Primer F, for amplifying $P_{gelE}$ from WT OG1RF. 15 bp overlap for InFusion cloning is underlined. |
| | oFR105 | CGTCAATGCCGTCATCAAACAATTAACTCCTTCCC | Primer R, for amplifying $P_{gelE}$ from WT OG1RF. 15 bp overlap for InFusion cloning is underlined. |
| | oFR100 | ATGCATGCTCGAGCATGGCAACTATCACAGATATC | Primer F, for amplifying GISE cassette from pSD15::$P_{gelE}$-DasherGFP. 14 bp overlap for InFusion cloning is underlined. |
| | oFR101 | GTACCGAGCTCGGATCATGACGTGTATTTCATTATCGA | Primer R, for amplifying GISE cassette from pSD15::$P_{gelE}$-DasherGFP. 16 bp overlap for InFusion cloning is underlined. |
| | oFR7 | GCTCGAGCATGCATCTAGAGG | Primer F, for inverse PCR (linearisation) of pGCP213 for GISE cassette insertion. 14 bp overlap for InFusion cloning is underlined. |
| | oFR8 | GATCCGAGCTCGGTACCAAG | Primer R, for inverse PCR (linearisation) of pGCP213 for GISE cassette insertion. 16 bp overlap for InFusion cloning is underlined. Also used as sequencing primer F for P-*pheS** insertion. |
| | oFR39 | AACCTGTCGTGCCAGCTG | Primer F, for inverse PCR (linearisation) of pFR212 for P-*pheS** insertion. 15 bp overlap for InFusion cloning is underlined. |
| | oFR40 | TCCCGACTGGAAAGCGG | Primer R, for inverse PCR (linearisation) of pFR212 for P-*pheS** insertion. 15 bp overlap for InFusion cloning is underlined. |
| | oFR41 | CTGGCACGACAGGTTCTAAGCTTGATTTTCGTTCGTG | Primer F, for amplifying P-*pheS** from pRV1. 15 bp overlap for InFusion cloning is underlined. |
| | oFR42 | GCTTTCCAGTCGGGACTGTCCGCTAATTCTTGCG | Primer R, for amplifying P-*pheS** from pRV1. 15 bp overlap for InFusion cloning is underlined. |
| | oFR10 | GGAAATGATTTACCTACTGCG | Sequencing primer F for inserts between OG1RF_11778 and OG1RF_11779. |
| | oFR11 | TTGCAATGCGATTGACG | Sequencing primer R for inserts between OG1RF_11778 and OG1RF_11779. |

| | | | |
|---|---|---|---|
| | M13F_ pUC (-40) | GTTTTCCCAGTCACGAC | Universal sequencing primer F for pSD15/pGCP213 inserts. |
| | M13R_ pUC (-26) | GTCATAGCTGTTTCCTG | Universal sequencing primer R for pSD15/pGCP213 inserts. |
| | oFR8 | GATCCGAGCTCGGTACCAAG | Sequencing primer F for P-*pheS** insertion. |
| | oFR14 | CTAGTGTAGCCGTAGTTAGGC | Sequencing primer R for P-*pheS** insertion. |
| Creation of OG1RFC/ OG1RFS | oFR102 | CTATTTTAGATGAATTGAGCTC GCGG | Primer F, for inverse PCR (linearisation) of pFR212 for $P_{gelE}$-DasherGFP replacement. 15 bp overlap for InFusion cloning is underlined. |
| | oFR118 | ATCAGTAAGCATGCGTGGTAC | Primer R, for inverse PCR (linearisation) of pFR212 for $P_{gelE}$-DasherGFP replacement. 15 bp overlap for InFusion cloning is underlined. |
| | oFR116 | ATTCATCTAAAATAGGTCACTA GTAAAGCGAACGAAAAAC | Primer F, for amplifying *cat* from EfaMarTn cassette. 15 bp overlap for InFusion cloning is underlined. |
| | oFR117 | CGCATGCTTACTGATAGAATGC GTGTGCTCTGC | Primer R, for amplifying *cat* from EfaMarTn cassette. 15 bp overlap for InFusion cloning is underlined. |
| | oFR127 | ATTCATCTAAAATAGGAAAAAA TCGCTATAATGACCC | Primer F, for amplifying *spc* from pBSU101::DasherGFP. 15 bp overlap for InFusion cloning is underlined. |
| | oFR128 | CGCATGCTTACTGATTTCAATA GTTACAAATTGTTTCAC | Primer R, for amplifying *spc* from pBSU101::DasherGFP. 15 bp overlap for InFusion cloning is underlined. |
| | oFR10 | GGAAATGATTTACCTACTGCG | Sequencing primer F for inserts between OG1RF_11778 and OG1RF_11779. |
| | oFR11 | TTGCAATGCGATTGACG | Sequencing primer R for inserts between OG1RF_11778 and OG1RF_11779. |
| Creation of OG1RFC Δ*gelE* | oFR94 | GATGCATGCTCGAGCGAATTGA AAATGTTCGCTATCTC | Primer 1, for amplifying 519 bp upstream sequence of *gelE* from WT OG1RF. 15 bp overlap for InFusion cloning is underlined. |
| | oFR129 | TCATTCATTGACCAGAACAGAC TTCATCAAACAATTAACTCCT | Primer 2, for amplifying 519 bp upstream sequence of *gelE* from WT OG1RF and ligation to downstream sequence. Complementary to oFR130. |
| | oFR130 | AGGAGTTAATTGTTTGATGAAG TCTGTTCTGGTCAATGAATGA | Primer 2, for amplifying 526 bp downstream sequence of *gelE* from WT OG1RF and ligation to upstream sequence. Complementary to oFR129. |
| | oFR98 | TACCGAGCTCGGATCGACGATC GTTTTGTTTGC | Primer 1, for amplifying 526 bp downstream sequence of *gelE* from WT OG1RF. 15 bp overlap for InFusion cloning is underlined. |
| | oFR7 | GCTCGAGCATGCATCTAGAGG | Primer F, for inverse PCR (linearisation) of pGCP213 to insert *gelE* deletion cassette. 15 bp overlap for InFusion cloning is underlined. |
| | oFR8 | GATCCGAGCTCGGTACCAAG | Primer R, for inverse PCR (linearisation) of pGCP213 to insert *gelE* deletion cassette. 15 bp overlap for InFusion cloning is underlined. |

| | | | |
|---|---|---|---|
| | M13F_ pUC (-40) | GTTTTCCCAGTCACGAC | Universal sequencing primer F for pFR213 inserts. |
| | M13R_ pUC (-26) | GTCATAGCTGTTTCCTG | Universal sequencing primer R for pFR213 inserts. |
| | oFR109 | GCAACAAATATTTACGCAGGGA AAG | Sequencing primer F for colony PCR of correct *gelE* deletion clones. |
| | oFR110 | GACAGGCTAAAACCGGCTATC | Sequencing primer R for colony PCR of correct *gelE* deletion clones. |
| Quantitative PCR (qPCR) | fsrA_F | AGCAACCTCAAATCCTGCCT | qPCR primer F for *fsrA* |
| | fsrA_R | TACAAGTGGCACACCAGGAC | qPCR primer R for *fsrA* |
| | fsrB_F | AGACCTTGGATGACGAGACCG | qPCR primer F for *fsrB* |
| | fsrB_R | GGTATGCGCCACAAGGAACA | qPCR primer R for *fsrB* |
| | fsrC_F | TGCACTGTTTTCAATCGCGT | qPCR primer F for *fsrC* |
| | fsrC_R | ACCGCAAAGCAAGCAAAACT | qPCR primer R for *fsrC* |
| | gelE_F | ACAAGATGGGCATCCCTCGA | qPCR primer F for *gelE* |
| | gelE_R | TCAAGCGCCATCACTAGCGA | qPCR primer R for *gelE* |
| | sprE_F | ATTGCGGTAGTGACTGTCGG | qPCR primer F for *sprE* |
| | sprE_R | CGACCATTGCGTGTGGTTTT | qPCR primer R for *sprE* |
| | entV_F | AGCTGCACAAAAGAAAGCCTG | qPCR primer F for *entV* |
| | entV_R | TAGCCCACATTGAACTGCCC | qPCR primer R for *entV* |
| | recA_F | GCGGCTGTTCCACCATTTCG | qPCR primer F for *recA* (housekeeping gene) |
| | recA_R | GTTGCATTGGGCGTAGGTGG | qPCR primer R for *recA* (housekeeping gene) |
